# Visualizing lipid nanoparticle trafficking for mRNA vaccine delivery in non-human primates

**DOI:** 10.1101/2024.06.21.600088

**Authors:** Maureen Buckley, Mariluz Araínga, Laura Maiorino, Ivan S. Pires, B.J. Kim, Katarzyna Kaczmarek Michaels, Jonathan Dye, Kashif Qureshi, Yiming Zhang, Howard Mak, Jon M. Steichen, William R. Schief, Francois Villinger, Darrell J Irvine

**Affiliations:** Department of Biological Engineering, Massachusetts Institute of Technology; Cambridge, MA, USA; Koch Institute for Integrative Cancer Research, Massachusetts Institute of Technology; Cambridge, MA, USA; Department of Chemical Engineering, Massachusetts Institute of Technology; Cambridge, MA, USA; Ragon Institute of Massachusetts General Hospital, Massachusetts Institute of Technology, and Harvard University; Cambridge, MA, USA; Howard Hughes Medical Institute; Chevy Chase, MD, USA; Department of Materials Science of Engineering; Massachusetts Institute of Technology; Cambridge, MA, USA; New Iberia Research Center, University of Louisiana at Lafayette, LA, USA; Consortium for HIV/AIDS Vaccine Development (CHAVD), The Scripps Research Institute, La Jolla, CA 92037, USA; Department of Immunology and Microbiology, The Scripps Research Institute, La Jolla, CA 92037, USA; IAVI Neutralizing Antibody Center, The Scripps Research Institute, La Jolla, CA 92037, USA

**Keywords:** mRNA vaccines, lipid nanoparticles, vaccine biodistribution

## Abstract

mRNA delivered using lipid nanoparticles (LNPs) has become an important subunit vaccine modality, but mechanisms of action for mRNA vaccines remain incompletely understood. Here, we synthesized a metal chelator-lipid conjugate enabling positron emission tomography (PET) tracer labeling of LNP/mRNA vaccines for quantitative visualization of vaccine trafficking in live non-human primates (NHPs). Following i.m. injection, we observed LNPs distributing through injected muscle tissue, simultaneous with rapid trafficking to draining lymph nodes (dLNs). Deltoid injection of LNPs mimicking human vaccine administration led to stochastic LNP delivery to 3 different sets of dLNs. LNP uptake in dLNs was confirmed by histology, and cellular analysis of tissues via flow cytometry identified antigen-presenting cells as the primary cell type responsible for early LNP uptake and mRNA translation. These results provide insights into the biodistribution of mRNA vaccines administered at clinically relevant doses, injection volumes, and injection sites in an important large animal model for vaccine development.

## INTRODUCTION

Major technological advancements for the use of mRNA as a therapeutic have been made over the past 20 years. Important innovations include the discovery of base modifications to modulate the lifetime and innate immune stimulatory capacity of mRNA, and the development of efficacious delivery vehicles that allow for effective delivery of mRNA *in vivo*^1–4^. For application in vaccines, pseudouridine base modifications weaken the recognition of mRNA by innate immune sensors such as endosomal Toll-like receptors and cytosolic RNA sensors, providing more efficient translation and antigen expression without induction of excess inflammation^2–3^. mRNA vaccines provide for a faster synthesis process than traditional protein-based vaccines and are readily manufactured at scale, making them a cost-effective solution for rapid therapeutic development^1^. During the SARS-COV-2 pandemic, the first FDA-approved mRNA vaccines were developed at record speed and proved to be highly effective in mitigating the incidence and severity of COVID-19^5^. Amongst different types of delivery vehicles, lipid nanoparticles (LNPs) stand as the most clinically advanced, with all currently approved mRNA vaccines utilizing LNPs to deliver their nucleic acid payloads^1,4^.

Although vaccines are most often administered intramuscularly, primary immune responses are initiated in draining lymph nodes. For traditional subunit vaccines, injected antigen is transported to draining lymph nodes (dLNs) via convection into draining lymph vessels or is taken up by antigen presenting cells that migrate from the injection site to dLNs^6^. In the lymph node, T cells and B cells undergo early activation, and a proportion of antigen-specific lymphocytes enter B cell follicles to form germinal centers^7–8^. Germinal centers (GCs) are critical immune hubs within the lymph node, working as the site of B cell affinity maturation and antibody diversification^7,9^. Antigen availability is critical to drive ongoing T and B cell responses. Though antigen in subunit vaccines is often rapidly cleared from lymph nodes following a primary immunization^10–12^, factors such as formulation of antigen in a nanoparticle form and immunogen glycosylation can impact antigen dispersal in T cell areas and interfollicular regions or promote antigen accumulation on follicular dendritic cells^12–16^. Such trafficking mechanisms have been interrogated for many different protein vaccine antigens through conventional staining and imaging modalities^12–13^.

While mechanisms of antigen dispersal and the biodistribution behavior of traditional protein vaccines combined with a variety of adjuvants have been studied in detail, much less is known about the fate of mRNA vaccines, especially in large animals and humans. Given its reliance on entry into cells for translation and the subsequent need for use of a delivery vehicle (most often, LNPs), the function of mRNA vaccines will be heavily influenced by the fate of the LNP. In mice, LNPs have been found to be taken up by conventional DCs and infiltrating immune cells in the lymph node, with mRNA vaccine-encoded reporter protein detectable in the subcapsular sinuses of lymph nodes^18–20^. In non-human primates (NHPs), mRNA/LNP vaccines were shown to induce strong innate immune activation in draining lymph nodes, with tagged mRNA payload and reporter protein-encoded mRNA detected at the injection site and dLNs following i.m. injection^21–22^. However, in these studies the specific muscle site that was injected was either not identified or multiple muscle injection sites were pooled for analysis, and thus it remains unclear if different anatomic sites (e.g., quadriceps vs. deltoid) exhibit different patterns of mRNA vaccine distribution. Notably, at very early times post immunization (4 hr), neither LNPs nor mRNA-expressed protein were detected in dLNs^21^, but both LNPs and mRNA payloads were readily measured in dLN antigen presenting cells (APCs) by 24 hr^21–22^. By contrast, a study of lipoplexes formed by complexation of mRNA with an aminoglycoside lipid CholK administered in the quadriceps of macaques and imaged by positron emission tomography (PET)/computed tomography (CT) whole-animal imaging detected RNA trafficking to multiple lymph nodes distal to the injection site within 4 hr^23^. The lipoplexes in this PET study have very different particle size, charge, and surface chemistry than the LNPs approved for use in humans, and thus it remains unclear if the rapid transport to LNs observed with this study is relevant for human mRNA/LNP vaccines. In humans, vaccine mRNA has been detected in biopsied axillary lymph nodes and blood of healthy volunteers, and at low levels in heart tissue but not liver or spleen of recently deceased vaccinees by RNA FISH (fluorescence *in-situ* hybridization) and qPCR^24–25^. At the protein level, vaccine antigens have been transiently detected at low levels in the blood of human vaccinees, and also localized in germinal centers of dLNs following booster immunization with the mRNA COVID vaccines^26^.

From this prior work, it has remained unclear (1) how reliable or stochastic is LNP uptake in different draining lymph node basins in large animals, (2) whether there is a significant contribution of direct drainage of LNPs to lymph nodes (vs. cell-mediated transport of mRNA/LNPs from the injection site), and (3) how many LNs are accessed following mRNA vaccination using clinically-relevant injection volumes/sites. To begin to address these questions, here we carried out studies tracking LNPs and mRNA in mice and non-human primates following administration of mRNA vaccines encoding reporter proteins or a stabilized HIV Env immunogen, using LNP compositions mimicking that used in the approved Moderna COVID vaccine. We developed radiometal chelator- tagged LNPs enabling whole-animal positron emission tomography (PET) imaging of the fate of vaccine carriers, and complemented whole-animal imaging of LNP distribution with tissue level histology, flow cytometry, and qPCR analysis of vaccine distribution in draining lymph node tissues. These studies revealed several facets of mRNA vaccine behavior in NHPs. Following i.m. injection, LNPs were localized in injected muscles, but could also be detected within 4 hr in draining lymph nodes, with stochasticity in terms of which lymph nodes accumulated vaccine. Combined immunofluorescence, qPCR and flow cytometry analyses on lymph nodes recovered 40 hr post immunization confirmed the presence of LNPs and mRNA in dLNs, and revealed antigen presenting cell (APC) populations as the primary cell types exhibiting LNP uptake and mRNA translation. These findings provide new insights into the biodistribution of mRNA vaccines at the whole-animal level and provide a rationale for understanding the potency of this vaccine modality.

## RESULTS

### Radiometal-chelating LNPs enable loading of PET tracers while retaining mRNA delivery function

To visualize LNP trafficking *in vivo*, we synthesized a lipid functionalized with the metal chelator 1,4,7,10-tetraazacyclododecane-1,4,7,10-tetraacetic acid (DOTA) that could be incorporated into the nanoparticles. Bicyclononyne-DOTA was reacted with aziodethyl phosphatidyl choline to form DSPC-DOTA (**Figure S1A**). The DOTA conjugate was purified by HPLC, and its identity verified by mass spectrometry (**Figure S1B-C**, **Table S1**). In parallel, base-modified mRNA encoding either mCherry as a reporter gene or a transmembrane form of the stabilized HIV Env trimer N332-GT2gp151 (note that we will refer to it as N332-GT2 for simplicity)^9,27^ was prepared by *in vitro* transcription. mRNA-loaded LNPs with compositions mimicking the Moderna COVID-19 mRNA vaccine formulation were prepared incorporating 0.5 mol% of the DSPC-DOTA (DOTA-LNPs). Dynamic light scattering (DLS) showed that LNPs prepared with the DOTA-lipid had slightly larger mean particle size, but similar overall particle size distributions, polydispersities, and zeta potentials as LNPs prepared without the DOTA lipid (**Figure 1A-B**). CryoTEM imaging of the LNPs also revealed similar morphologies for the LNPs prepared with or without incorporated DSPC-DOTA (**Figure 1C**). To confirm that inclusion of the chelator lipid did not affect the transfection activity of the LNPs, C2C12 murine myoblast cells were transfected with mCherry mRNA encapsulated in non-tagged LNPs or DOTA-LNPs loaded with ^64^Cu that had been allowed to decay to undetectable levels of radioactivity. Transfection efficiencies and mean fluorescence intensities of mCherry expression were identical between the two groups (**Figure 1D-F)**. Thus, incorporation of a low level of tagged lipid enables LNPs to be generated that carry a radiometal chelator and are fully functional for mRNA delivery.

**Figure 1.**
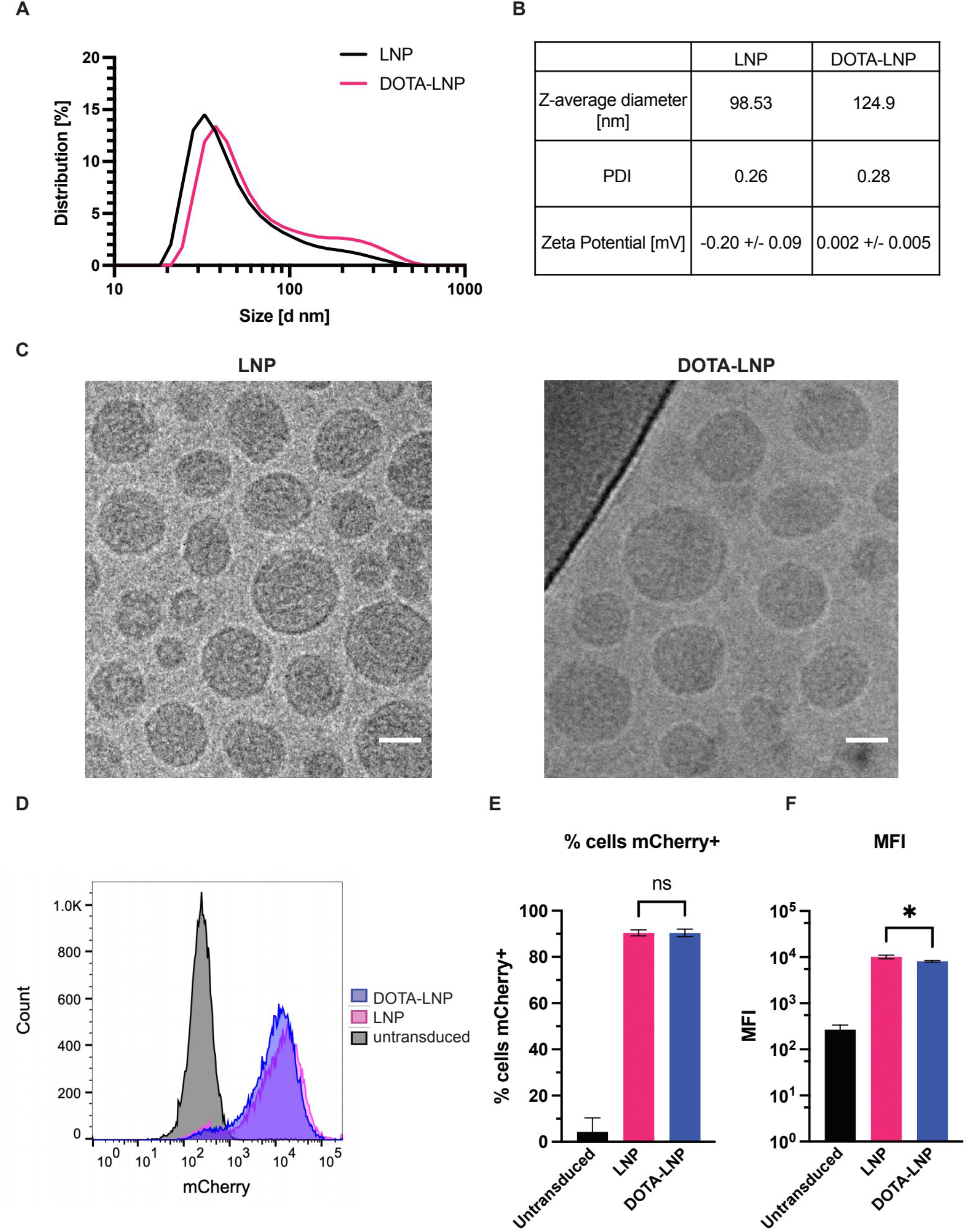
Radiometal-chelating LNPs enable loading of PET tracers while retaining mRNA delivery function. **(A)** dynamic light scattering (DLS) analysis of LNPs with or without DOTA-lipid. **(B)** LNP quality control metrics table for LNPs with or without DOTA- lipid. **(C)** CryoTEM imaging of LNPs and DOTA-LNPs. Scale bars 50 nm. **(D-F)** C2C12 cells were incubated with 10 µg/mL mCherry-encoding mRNA delivered by LNPs or ^64^Cu-loaded DOTA-LNPs for 24 hrs, then analyzed by flow cytometry for mCherry expression. Shown are histograms of mCherry fluorescence (C), percentages of mCherry positive cells (D), and mean fluorescence intensities of transfected cells (E). Statistical significance was determined by one-way ANOVA followed by Tukey’s post hoc test. *P < 0.05; ****P < 0.0001. All data show means ± SEM.

### LNPs distribute in injected muscle and rapidly reach draining lymph nodes in mice

To evaluate radiometal loading, ^64^Cu was added to DOTA-LNPs for 60 min followed by dialysis to remove unbound copper. Thin layer chromatography analysis showed effective loading of the LNPs with ^64^Cu and removal of free metal (**Figure S2A-C**). To validate the utility of ^64^Cu/DOTA-labeling for assessing the biodistribution of LNP- mRNA, BALB/c mice were injected with 5 ug mCherry mRNA encapsulated in ^64^Cu-loaded DOTA-LNPs **(Figure S2D)** i.m. in the right gastrocnemius muscle. As a control, a separate cohort of mice were injected with free ^64^Cu (same activity and volume as DOTA-LNPs) to distinguish the behavior of free ^64^Cu. Animals were imaged via whole body PET-computed tomography (CT) at 0h, 3hr, 6hr, and 24hr post immunization; at 24 hr tissues were isolated for *ex vivo* PET imaging (**Figure 2A**). Labeled LNPs were clearly visible dispersing in the injected muscle immediately following injection (**Figures 2B** and **S3**). Further, by 3 hr signals could be detected in multiple draining lymph nodes, and for some animals, LNPs were detected at the needle entry point in the muscle (suggesting some leakage along the needle track, **Figure 2B**). Popliteal LNs showed LNP signal in all of the mice, while stochastic uptake in inguinal, iliac, or more distal axillary LNs was observed in some animals (**Figures 2B** and **S3**, **Videos S2-S4**). Free copper by contrast showed rapid clearance from the injection site and no accumulation in draining nodes (**Figure S3**, **Video S1**). We quantified mean standard uptake values (SUVmean; PET signal normalized to dose and body weight) at the injection sites, draining popliteal LNs, and liver over time. Free copper cleared from the injection site almost entirely within 3 hr, while DOTA-LNPs showed a slower decay, with ∼40% of the LNP signal initially deposited still present at 24hr (**Figure 2C**). Free copper showed no accumulation in draining LNs, while DOTA-LNP signal accumulated by 3 hr in all animals, and was still present at 24 hr (**Figure 2D)**. Such a rapid transport to LNs suggests direct drainage of LNPs via lymphatics as it is too rapid to reflect cell-mediated transport. Signal from DOTA-LNPs in the liver was also seen to increase with time, but was an order of magnitude lower than signal detected in draining LNs and substantially lower than signal from the free ^64^Cu control (**Figure 2E**).

**Figure 2.**
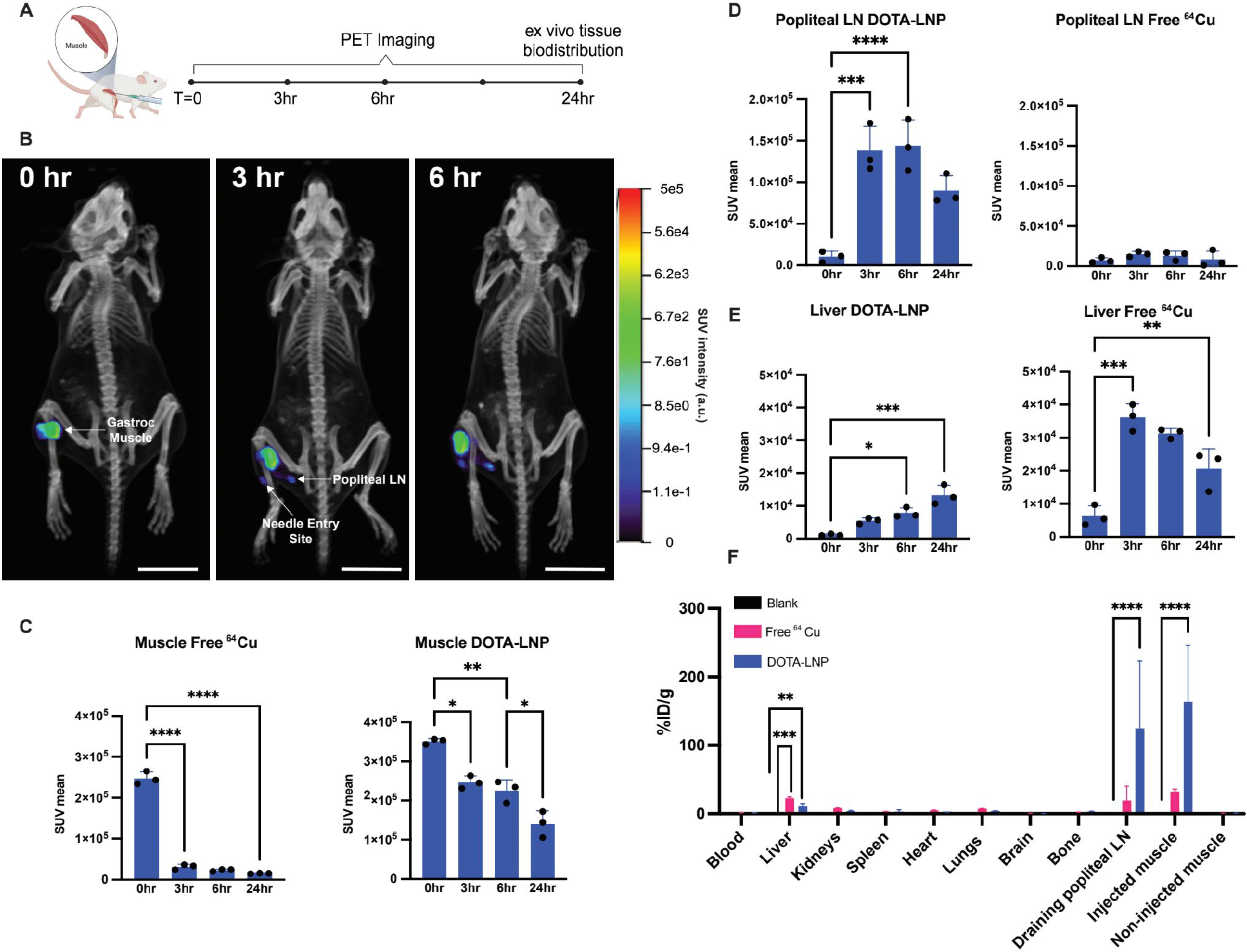
PET-CT imaging reveals LNPs primarily distribute at injected muscle at immediate draining lymph node in mice. (A) PET-CT study timeline. DOTA-LNPs encapsulating 5 µg mCherry mRNA were administered i.m. into the gastroc muscle of BALB/c mice (*n* = 3 animals/group). **(B)** PET-CT projections of mice over the imaging time course. Scale bars 50 mm. **(C-E)** ROI analyses of PET signal at injection site (gastrocnemius muscle, C), draining popliteal LNs (D), and liver (E) for animals receiving free ^64^Cu or ^64^Cu-loaded DOTA-LNPs. **(F)** E*x vivo* tissue gamma counter measurements of ^64^Cu signal from DOTA-LNPs compared to blank and free ^64^Cu controls. Statistical significance was determined by two-way ANOVA followed by Tukey’s post hoc test. *P < 0.05; **P < 0.01; ***P < 0.001; ****P < 0.0001. All data show means ± SEM.

The DOTA-LNP-injected animals were euthanized at 24 hr and organs were harvested for *ex vivo* activity quantification via gamma counter. The only significant differences in signal between the blank control and DOTA-LNPs were seen in injected muscle and popliteal draining lymph node tissues, with minor liver signal remaining, which was lower than that of Free ^64^Cu **(Figure 2F)**. Overall, PET-CT and gamma quantification analysis showed a strong trafficking preference for LNPs post-i.m. injection to distribute in the injected muscle and proximal draining lymph nodes, with relatively high retention of LNPs exhibited at the injection site, which was distinct from the behavior of free copper.

### LNPs distribute between the injection site and draining lymph nodes in rhesus macaques

We next evaluated the biodistribution of mRNA-loaded LNPs in rhesus macaques. DOTA-LNPs were loaded with radioactive ^64^Cu as before and mixed 1:1 with LNPs labeled with the lipophilic fluorophore DiD, to enable whole-animal PET imaging followed by tissue-level biodistribution/imaging analyses. Both LNPs encapsulated mRNA either encoding for mCherry fluorescent protein or a transmembrane form of the stabilized HIV Env trimer N332-GT2^12^. In a first set of 4 animals, LNPs carrying mCherry mRNA were injected in the left deltoid and right quadriceps (50 µg mRNA per injection site), with animals imaged via whole body PET-CT at 0 hr, 4 hr, and 24 hr **(Figure 3A**, left schematic, **Videos S5-S7)**. In a second cohort of 4 animals, LNPs encapsulating N332-GT2 mRNA were administered only in the left deltoid, and followed the same PET-CT imaging timeline (**Figure 3A**, right schematic, **Videos S8-S10**).

**Figure 3.**
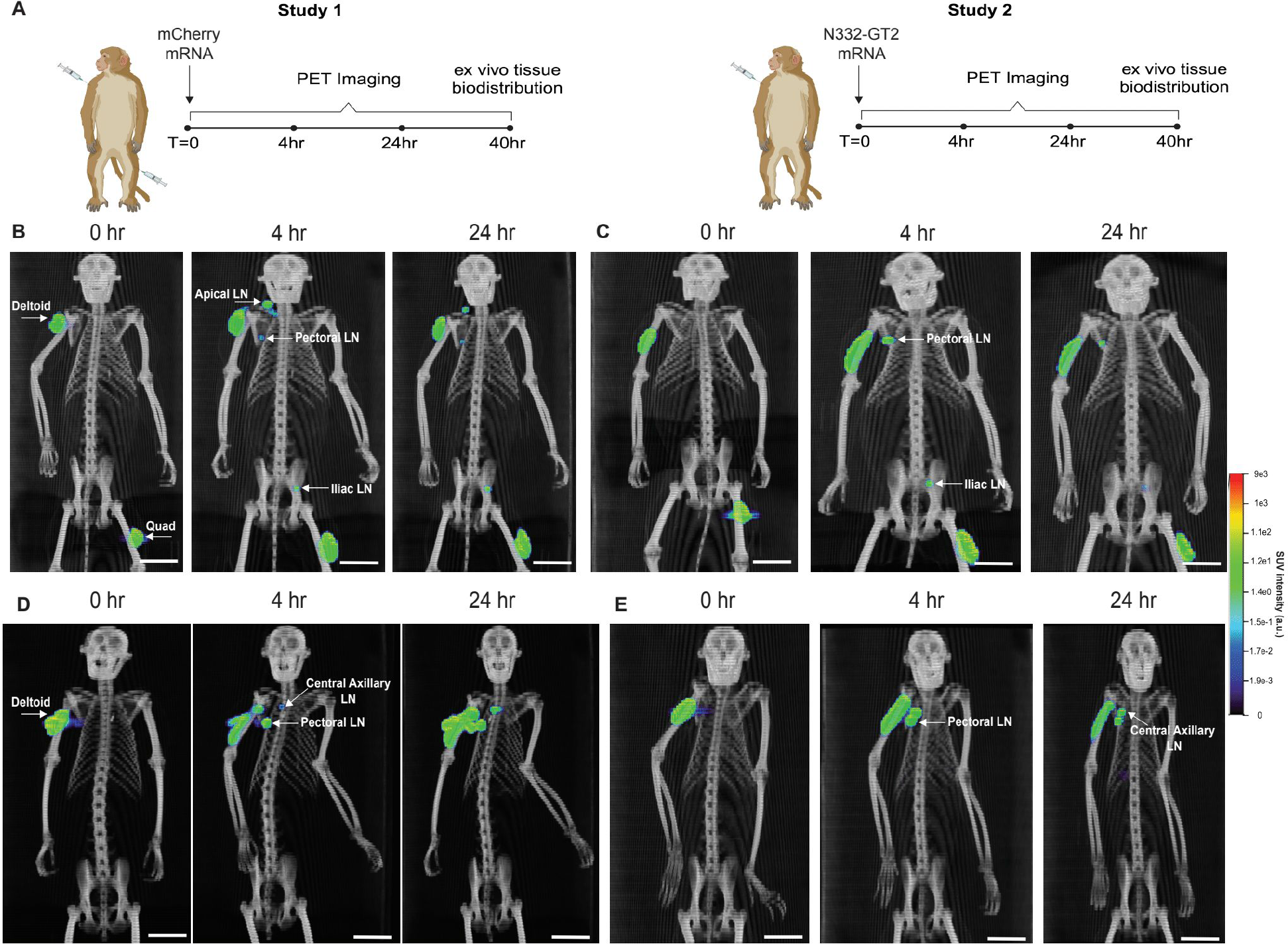
LNPs rapidly reach draining lymph nodes following i.m. administration in rhesus macaques. (A) Schematics of PET-CT timelines for mCherry (left) and N332- GT2 (right) NHP studies. Animals received injections of 50 µg mRNA per site i.m. **(B)** PET-CT projections of one representative animal from the mCherry study over time. **(C)** PET-CT projections of a second representative animal from the mCherry study over time. **(D)** PET-CT projections of one representative animal from the N332-GT2 study over time. **(E)** PET-CT projections of a second representative animal from the mCherry study over time. Scale bars in B-E represent 50 mm.

The whole-animal images revealed distinct draining LN basins for the deltoid vs. quadriceps injections. For the quadriceps injection site, at 4 hr, LNP signal was distributed through the injected muscle, and could also be clearly detected in iliac LNs (**Figure 3B-C**, **Figure S4**, and **Video S6)**. LNPs injected i.m. in the deltoid by contrast drained stochastically to axillary, apical, or pectoral lymph nodes, with different LNs exhibiting LNP uptake in different animals, **(Figure 3B-E, Figure S5,** and **Videos S6, S9)**. Although multiple lymph nodes may reside at each of these drainage sites, the resolution of the PET scans did not permit clear identification of whether one or more nodes took up vaccine at any of the individual drainage sites. Low signal was also detected in the spleen of 2 NHPs but was negligible in the majority of animals (**Figure S5**). LNP signals were not detected in the heart or other tissues.

As the deltoid LNP biodistributions were similar in the two cohorts, we pooled the quantitative analyses of the muscle and dLNs for this injection site. Decay-corrected PET signals in the injected muscle sites steadily decreased from 0 to 24 hr, suggesting dissemination of LNPs from the injection site (**Figure 4A-B**). Draining lymph nodes by contrast showed peak signal at 4 hr, followed by a significant drop in LNP signal by 24 hr **(Figure 4C-D)**. Annotation of LNP uptake animal by animal showed that iliac LNs were preferentially targeted following quadriceps injection, and we only detected signals in single lymph node sites (**Figure 4E**). From the deltoid injections, LNP uptake was stochastically detected in axillary, apical, and pectoral LNs, and a majority of animals (5 out of 8) showed uptake in 2 different dLN sites, while 1 of 8 animals showed no uptake in draining lymph nodes over the time course (**Figure 4F**). Consistent with our findings in mice, LNP signal in the liver was detectable but ∼10-fold lower than that detected in draining lymph nodes; and the signal detected in the spleen and heart was also very low (**Figure 4G-I)**.

**Figure 4.**
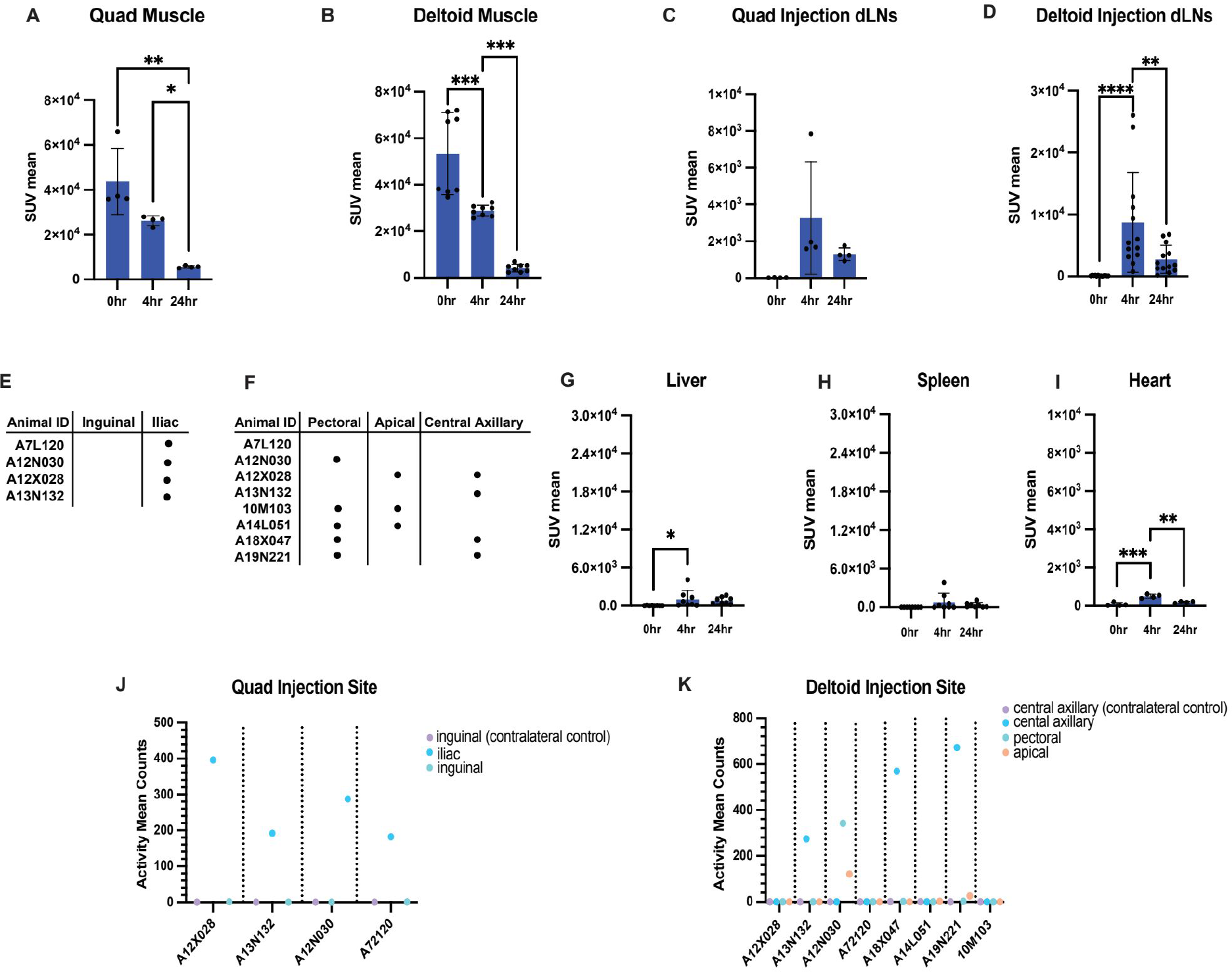
LNPs access draining lymph nodes but show very limited systemic distribution following i.m. injection in rhesus macaques. (A-D) ROI analyses of PET signal at quadriceps muscle injection site (A), deltoid muscle injection site (B), quadriceps draining lymph nodes (C), and deltoid draining lymph nodes (D). **(E-F)** Individual animal tables marking lymph nodes with detectable signal in the quadriceps draining lymph node region (E) and the deltoid draining lymph node region (F). **(G-I)** ROI analyses of PET signal at liver (G), spleen (H), and heart (I). (J-K) *Ex vivo* PET activity reads for draining and non-draining lymph nodes at quadriceps injection site (J) and deltoid injection site (K). Statistical significance was determined by one-way ANOVA followed by Tukey’s post hoc test. ns, P > 0.05; *P < 0.05; **P < 0.01; ***P < 0.001; ****P < 0.0001. All data show means ± SEM.

At 40 hr post-injection, the animals were euthanized and select tissues were harvested for *ex vivo* PET-CT analysis. In quad-draining LNs, all animals exhibited LNP signal remaining at this time point in iliac LNs (**Figure 4J**). For the deltoid injections, 3 animals showed clearly detectable LNP signal in axillary LNs, while lower signal was seen in apical LNs in a few animals, and 4 of 8 animals showed no remaining detectable LNP signal in any of the nearby LNs by this timepoint injection **(Figure 4K)**. Thus, LNPs distribute primarily in the injected muscle and nearby draining lymph nodes early after mRNA immunization.

### Tissue-level analysis confirms LNPs and vaccine mRNA are delivered to multiple draining lymph nodes in NHPs

At 40 hr post injection, the 4 animals receiving LNPs carrying mCherry mRNA in the quadriceps were euthanized and draining iliac lymph nodes were harvested for tissue-level analysis. We first carried out confocal imaging of histological tissue slices to visualize the biodistribution of LNPs in quadriceps-draining iliac LNs. When compared to control contralateral lymph nodes, substantial LNP signal (green) was detected in the periphery of the tissues **(Figure 5A-B)**.

**Figure 5.**
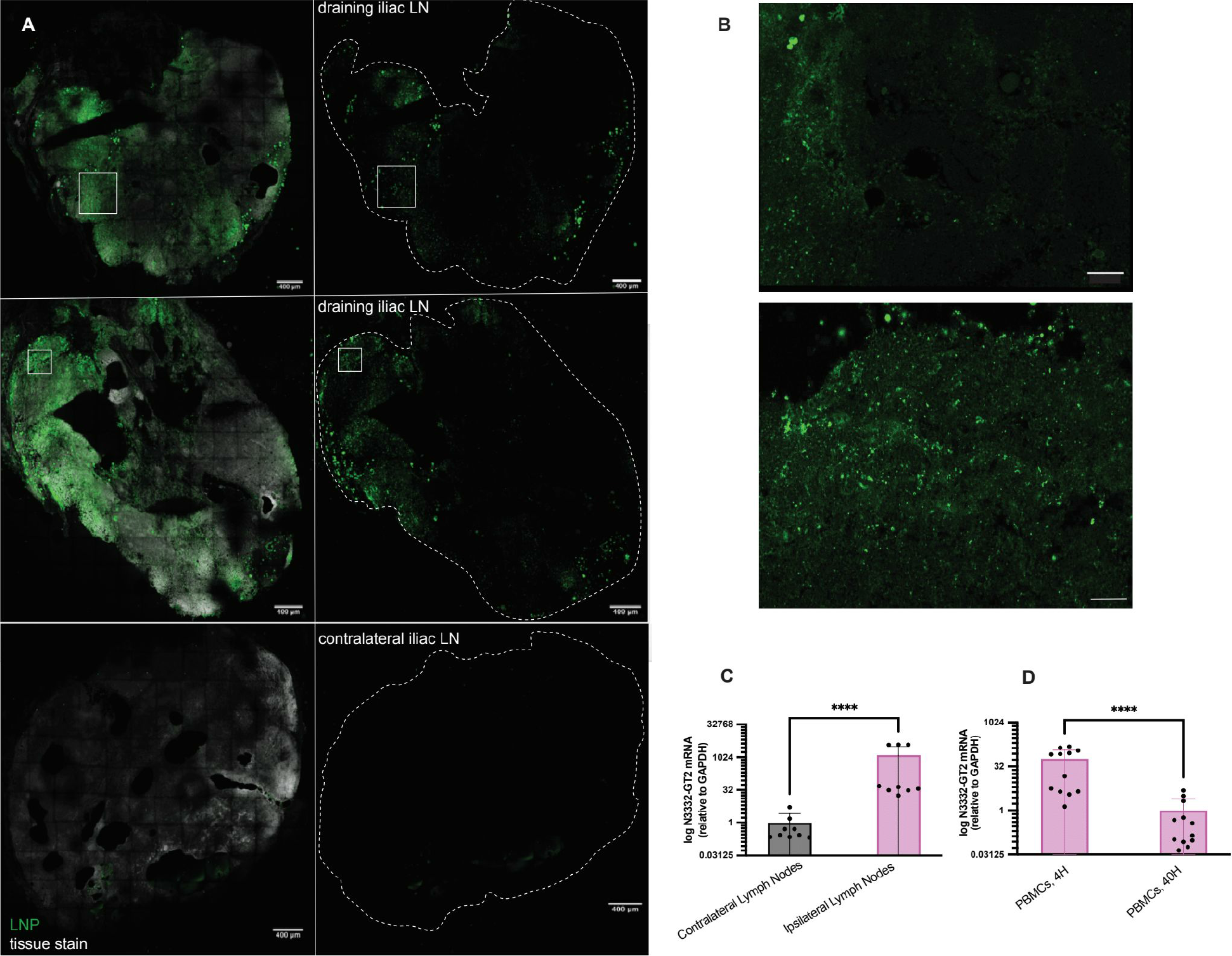
Ex vivo tissue analysis reveals LNPs and mRNA are persistent in draining lymph nodes of NHPs. (A) NHP draining and non-draining (contralateral) lymph node samples taken at 40h post-immunization. LNP signal via diD in *green*, tissue stain in *grey*. Magnification at 25x with top panel showing merged signal and bottom panel showing only LNP signal. Scale bars 400 um **(B)** 63x magnification comparing LNP signal (green) between selected draining (white box indicating zoomed in region of view in 25x image) and non-draining lymph node samples. Scale bars 115 um. **(C)** Expression of N332-GT2 mRNA in lymph nodes from ipsilateral (right) and contralateral (left) side measured by qRT-PCR using GAPDH as a reference gene. The experiment represents apical, central axillary, and pectoral lymph nodes obtained from single animal 24 hours post-injection. Three technical replicates were performed for each sample. **(D)** Expression of N332-GT2 mRNA in sorted PBMCs at 4 hours and 24 hours post-injection measured by qRT-PCR using GAPDH as a reference gene. These experiments represent PBMCs obtained from four animals. Three technical replicates were performed for each sample. Statistical significance was determined by Mann Whitney test. ns, P > 0.05; *P < 0.0332; **P < 0.0021; ***P < 0.0002; ****P < 0.0001. All data show means ±SEM.

To confirm that mRNA was also delivered to dLNs, we carried out qPCR analysis to detect Env trimer mRNA in deltoid-draining LNs collected from the second PET study. Trimer mRNA was detected in axillary and pectoral lymph nodes, with the greatest amount detected in the central axillary lymph nodes **(Figure 5C)**. In addition, qPCR was run on collected PBMCs (peripheral blood mononuclear cells) to assess mRNA in circulating cells at 4 hr and 40 hr post-injection. Trimer mRNA was detected in PBMCs from 3 of 4 animals at 4 hr post-immunization, but was undetectable at 40 hr **(Figure 5D)**. These findings corroborate the PET/CT imaging data suggesting LNP/mRNA reaches multiple draining lymph nodes at early time points post immunization.

### APCs found to be cell type responsible for LNP uptake and mRNA translation in draining LNs

Following PET-CT imaging of the 4 animals receiving LNPs carrying mCherry mRNA, draining and contralateral control lymph node tissues were harvested to investigate the cellular distribution of DOTA-LNPs and mRNA-encoded protein using flow cytometry **(Figures S6-7)**. Fluorescent LNPs were readily detected in a small proportion of lymph node cells collected from draining lymph nodes at both the deltoid and quadriceps injection sites (**Figure 6A-B**). Consistent with the PET-CT findings indicating a greater frequency of animals exhibiting LNP trafficking to iliac LNs, LNPs injected at the quadriceps were primarily detected in cells from iliac but not nearby inguinal nodes (**Figure 6A-B)**. From LNs analyzed at the deltoid injection site, LNP uptake was primarily detected in apical and pectoral LNs, but not the central axillary LN (**Figure 6A- B).** However, only two of the deltoid-draining LNs we analyzed showed a relatively high (>1%) LNP signal, limiting our ability to draw significant conclusions for drainage at this area. Although we expected mRNA expression to be substantially decayed by the time point these tissues were collected (40 hr post immunization), we also assessed mCherry signal in LN cells. Low but statistically significant mCherry expression was detected in cells from the quadriceps-draining iliac LNs, and in 2 apical LNs draining the deltoid immunization site (**Figure 6C-D**).

**Figure 6.**
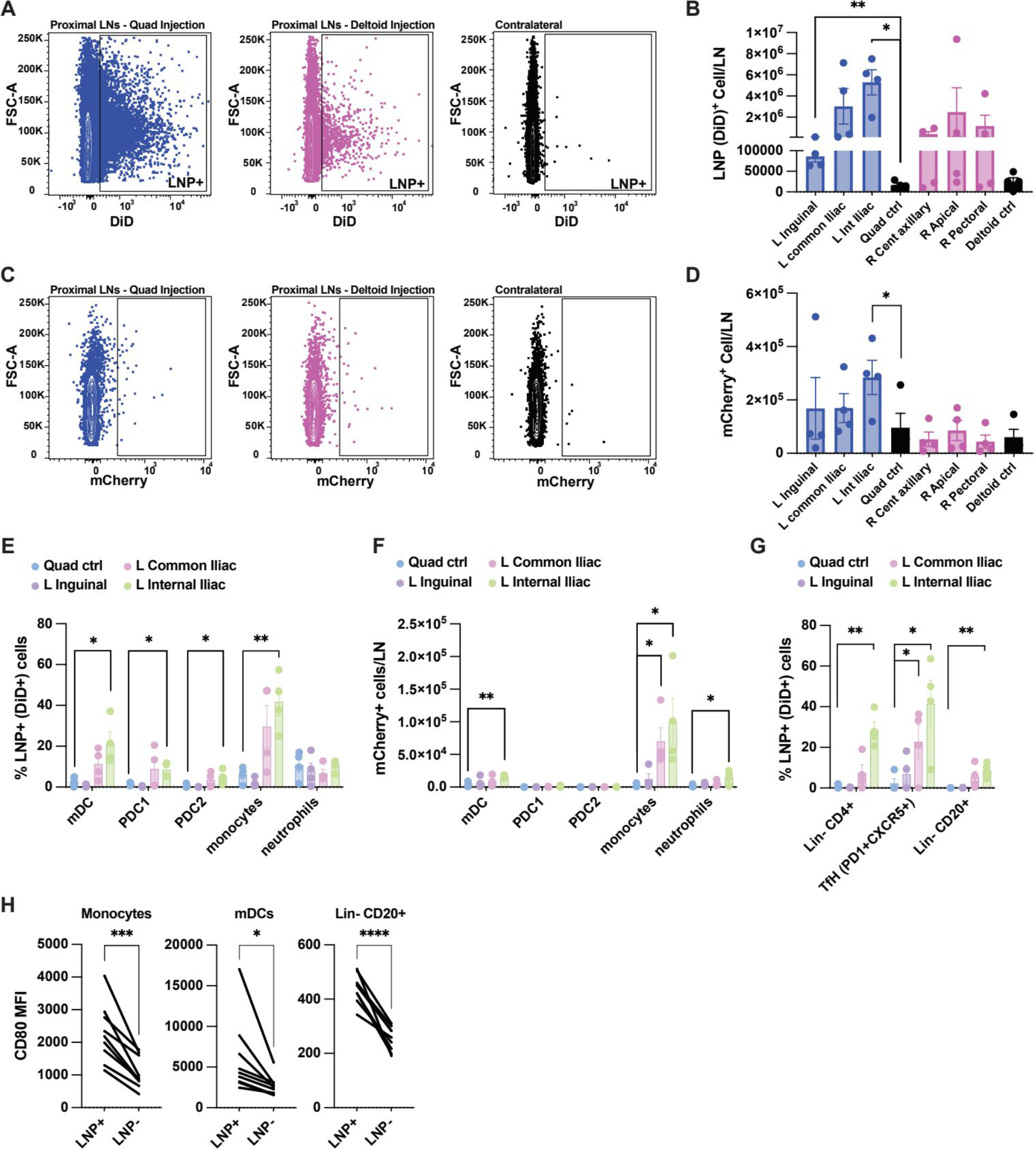
Monocytes and dendritic cells acquire LNPs in NHP draining LNs. (A) Representative flow cytometry contour plots of LNP (DiD) signal in all cells. **(B)** Total cell counts gated on LNP^+^ (DiD) signal. **(C)** Representative flow cytometry contour plots of mCherry signal. **(D)** Total cell counts gated on mCherry^+^ signal. **(E)** Percentage cells positive for LNP signal within specific myeloid populations in LNs draining the quadricep injection site. **(F)** Number of cells positive for mCherry^+^ signal within specific myeloid populations in LNs draining the quadricep injection site. **(G)** Percentage cells positive for LNP signal within specific lymphoid populations in LNs draining the quadricep injection site. **(H)** Geometric mean fluorescent intensity of CD80-BV650 signal in LNP^+^ relative to LNP^-^ cells within LNs with high LNP uptake (>1%). Cell populations were gated as follows: mDCs (SSC^high^CD14^-^CD16^-^ MHCII^+^CD11c^+^); pDC1s (SSC^low^MHCII^+^CD123^+^); pDC2s (SSC^high^CD14-CD16-MHCII^+^CD11c^-^CD123^+^); Monocytes (SSC^high^MHCII^+^CD14^+^ and/or CD16^+^); Neutrophils (SSC^high^MHCII^-^CD66abce^+^); Lin^-^(CD8^-^CD14^-^CD16^-^).

To examine which cell populations were taking up LNPs, we first assessed changes in myeloid cell populations in the dLNs that may have been induced as part of the innate immune response triggered by LNP-mRNA immunization^28–30^. Interestingly, in the quadriceps-draining LNs, mDCs and monocytes were significantly increased relative to the contralateral control nodes **(Figure S8A)**. There was a trend toward increased mDCs and monocytes detected in deltoid-draining LNs relative to contralateral control LNs, but these changes did not reach statistical significance (**Figure S8B**). We next examined the association of LNP and mCherry signal with different LN cell populations. In the quadriceps-draining LNs, myeloid DCs and monocytes showed the highest uptake of LNPs, and these were the cell types that also were detected with mCherry expression, along with a small population of neutrophils **(Figure 6E-F)**. LNP uptake was also detected in CD4^+^ T cells and B cells, but mCherry^+^ cell numbers were too low to draw any conclusions about expression in these cell types **(Figure 6G** and data not shown). An analogous analysis of the deltoid-draining LNs showed similar trends for LNP uptake in mDCs and monocytes, but these observations did not reach statistical significance (**Figure S8C-F**). Notably, we found that antigen presenting cells that were LNP^+^ also upregulated the activation marker CD80 **(Figure 6H)** and to a lesser extent CD40 and MHCII **(Figure S8G, H)** relative to LNP^-^ cells within the same LN. Altogether, this data indicates that a variety of cells with antigen presenting/phagocytic functions are the primary cells acquiring LNPs.

## DISCUSSION

mRNA delivered by lipid nanoparticles has emerged as an important clinical modality for vaccines, with millions of doses safely administered during the COVID pandemic. However, much of the underlying mechanisms of action for mRNA vaccines *in vivo* remain to be understood. Here we sought to gain insights into the early biodistribution and kinetics of LNP distribution in tissues in rhesus macaques, an important preclinical animal model for vaccine development and the closest animal model genetically to humans. Using whole-animal PET/CT imaging and testing two administration sites, we identified several features of LNP pharmacokinetics following intramuscular injection: First, LNPs spread within the injected muscle tissue immediately following injection, but also rapidly showed uptake in distinct draining lymph node pools. Second, the distribution of LNPs among different draining lymph node basins was stochastic animal to animal, particularly for deltoid i.m. injections, where three different lymph node regions– axillary, apical, and pectoral LNs- were variably involved animal to animal. Third, lymph node accumulation was observed within 3 hr post injection, a timespan suggesting that direct transport through lymphatics plays a role in LNP accumulation in dLNs. This accumulation of LNPs in lymph nodes observed by live animal imaging was corroborated by histological and flow cytometry analyses at later time points.

In preliminary studies in mice, we found, similar to our data in NHPs, LNPs rapidly distributed both in the injected muscle and proximal draining lymph nodes, with negligible signal in other tissues except for low levels of uptake in the liver. Previous studies investigating LNP biodistribution in mice and rats have identified the liver as a hot spot for LNP drainage and accumulation at early time points post- injection, following both i.v. and i.m. injections^31–32^. However, we found that liver uptake following i.m. injection is primarily promoted by injection of larger LNP dosages and/or volumes (preliminary results not shown), which leads to spillover of LNPs from the injection site into the blood^33^.

In macaques, the stochasticity in terms of which dLNs showed LNP uptake from the deltoid injection site appears to be a feature of the NHP model that may be distinct from humans, as longitudinal fine needle aspirate studies of volunteers receiving the COVID-19 mRNA vaccines have been quite successful in detecting robust ongoing germinal center responses simply by sampling draining axillary LNs^34,35^. Human autopsy studies completed on patients that had received the SARS-CoV-2 mRNA vaccine prior to passing identified the presence of vaccine mRNA in draining lymph nodes and some heart tissues using RT-qPCR^24^. We detected little or no LNP signal in heart tissue via PET imaging following i.m. immunization of NHPs. In alignment with what we observed in NHPs by PET, no vaccine mRNA was detected in the liver in humans^24^.

Rapid LNP delivery to draining lymph nodes was observed here in both mice and rhesus macaques, with the 4 hr timepoint exhibiting max LNP signal. This finding aligns with other work done to non-invasively track mRNA lipoplexes through radiolabeling of the mRNA payload, where uptake in distal para-aortic LNs from a quadriceps injection site was detected 4 hr post injection^23^. Such early signal detection in the draining lymph nodes is too rapid to reflect cell-mediated transport and is most consistent with direct transport of LNPs via lymph. Cell-mediated trafficking of LNPs/mRNA from the injection site could still occur over a period of multiple days post-immunization, which is beyond the timeframe that our ^64^Cu radiotracer could reliably report. We also saw decays in LN LNP signal between 4 and 24 hr, even though fluorescently-tagged LNPs were still readily detected by histology 40 hr post injection. This may suggest that the DOTA- tagged lipid used to track the LNPs here may be rapidly metabolized *in vivo*.

LNP biodistribution patterns seen by whole-animal PET imaging were corroborated by flow cytometry and histological imaging analysis of individual draining lymph nodes. mRNA vaccines have been reported to trigger an increase in myeloid cells in draining lymph nodes that are capable of taking up LNPs and initiating an innate immune response^21^. In accordance with these observations, we found myeloid cells to increase in mRNA vaccinated animals, with antigen-presenting cells primarily responsible for LNP uptake. Additionally, monocytes, neutrophils, and DCs were similarly identified as cells responsible for translating vaccine mRNA. These findings are largely in agreement with prior work that characterized LNP uptake and mRNA expression by flow cytometry in NHP lymph nodes at earlier time points (24 hr and earlier).^21–22^.

In summary, PET/CT imaging is a powerful modality to gain a whole-animal level view on the biodistribution of vaccines in non-human primates. Using metal chelator-conjugated lipids, we were able to visualize LNP/mRNA vaccine biodistribution in both mice and NHPs. These studies identified stochastic transport to proximal draining lymph nodes with limited distribution into other tissues. These data provide further insights into the key lymph nodes involved in the immune response to mRNA vaccines in the closest animal model to humans, and can guide further mechanistic studies of key draining lymph nodes where mRNA vaccines act.

## ACKNOWLEDGEMENTS

We thank the Koch Institute Swanson Biotechnology Center’s Flow Cytometry, Microscopy, Histology, PMIT, and Nanotechnology Materials core facilities for their technical support. This work was supported in part by the Koch Institute Support (core) Grant 5P30-CA014051 from the National Cancer Institute, the NIH (award AI176533 to DJI and award UM1AI144462 to WRS and DJI), and the Ragon Institute of MGH, MIT, and Harvard. We extend our thanks to Douglas Ferrell for assistance in PET/CT scanning of nonhuman primates and the staff at the New Iberia Research Center for their expert care of the nonhuman primates.

## AUTHOR CONTRIBUTIONS

M.B. performed DOTA-LNP formation and mRNA encapsulation, LNP QC protocols, C2C12 cell transfection, *in vitro* flow cytometry, i.m. injections and necropsies for mouse study, PET-CT data analysis, *in vivo* flow cytometry processing and analysis, tissue staining, imaging, and analysis for IF, and wrote the manuscript. L.M. performed NHP shipping and tissue handling, *in vivo* flow cytometry and qPCR processing and analysis, and edited the manuscript. I.S.P synthesized and validated DSPC-DOTA via HPLC and mass spectrometry. B.K. helped develop DOTA-LNP formulation and optimized QC protocols for LNPs. K.K.M performed qPCR RNA extraction and processing. J.D. produced mRNA constructs used. K.Q. performed qPCR RNA extraction and processing. H.M. performed ^64^Cu loading, mouse PET-CT imaging and necropsies, and *ex vivo* tissue biodistribution processing and analysis. M.A. performed PET imaging and necropsy. F.V. designed the experiments, and performed PET imaging, animal care and necropsies. J.M.S. and W.R.S. provided the N332-GT2 gp151 immunogen amino acid sequence. D.J.I. designed the experiments and edited the manuscript.

## Supplemental Information

Document S1. Figures S1-S8 and Table S1 Videos S1-S10

## STAR METHODS

### mRNA synthesis

Template DNA plasmids used in the production of mRNA were created using a commercially available Cloning Kit for mRNA Templates (Takara #6143) according to the manufacturer’s instructions. Resultant plasmid DNA was linearized via endonuclease digestion and purified with PureLink PCR Purification columns (ThermoFisher #K310002) following the manufacturer’s instructions. To synthesize RNA, 20 μL *in vitro* transcription (IVT) reactions were performed using reagents from the HiScribe T7 High Yield RNA Synthesis Kit (NEB #E2040) and 1-2 μg of linear DNA template (scaled as needed). Modified base N1-methylpseudouridine triphosphate (TriLink #N-1081) was added to the reaction mixture instead of canonical uridine triphosphate, and CleanCap Reagent AG (TriLink #N-7113) was utilized to co- enzymatically add 5’ Cap-1 structures to synthesized RNA. The IVT product was purified using PureLink RNA Mini columns (ThermoFisher #12183018A) following the manufacturer’s instructions. Quality of the resulting mRNA was assessed using UV-Vis spectrophotometry and gel electrophoresis.

### DSPC-DOTA synthesis

1,2-distearoyl-sn-glycero-3-phosphocholine (N-azidoethyl) (18:0 azidoethyl PC, Avanti Polar Lipids) was dissolved in chloroform at 10 mg/mL and reacted with a 3-fold molar excess of methanol-dissolved BCN-DOTA (CheMatech). The reaction was allowed to occur for at least 1 day at 4 °C with shaking at 5 mg/mL lipid in a 50:50 mixture of chloroform and methanol. After completion of the reaction, DSPC-DOTA was purified from excess BCN-DOTA via reverse-phase high-pressure liquid chromatography (RP-HPLC) on a Jupiter C4 column (5 µm particles, 300 Å – Phenomenex P/N: 00G-4167- E0) following the gradient shown in **Table S1**. Fractions corresponding to DSPC-DOTA were collected and methanol was removed on a rotovap. The sample was then diluted with a 10x volume of deionized water, frozen in liquid nitrogen and lyophilized.

### mRNA encapsulation in LNPs

Lipids were stored in ethanol at −20 °C, and RNA constructs were stored in RNAse-free water at −80 °C and were thawed on ice before use. The two phases were prepared at an ethanol:aqueous volume ratio of 1:2, and RNA and lipids combined at an N:P ratio of 5:1. Each phase was loaded into a syringe (BD), and locked onto the NxGen microfluidic cartridge for mixing using a NanoAssemblr Ignite instrument (Precision Nanosystems). The Ignite was set to operate with the following settings: volume ratio- 2:1; flow rate- 12 mL/min; waste volume- 0 mL.

The organic phase was prepared by solubilizing the lipids SM102 (BroadPharm CAT#25499), DSPC (Avanti Polar Lipids CAT#850365), Cholesterol (Avanti Polar Lipids CAT#700100), DMG-PEG2k (Avanti Polar Lipids CAT#88015), and DSPC-DOTA (see *DSPC-DOTA* synthesis) in ethanol at a molar ratio of 50:9.5:38.5:0.5:1.5 and a total lipid concentration of 5 mg/ml. For non-DOTA lipids, SM102, DSPC, Cholesterol, and DMG- PEG2k were solubilized in ethanol at a molar ratio of 50:10:38.5:1.5 and a total lipid concentration of 5 mg/ml. Non-DOTA lipids labeled with fluorescent diD were solubilized in ethanol at a molar ratio of 50:0.1:9.9:38.5:1.5 and a total lipid concentration of 5 mg/ml. The aqueous phase of RNA was prepared by diluting the RNA (stored in RNAse-free water) with 10 mM citrate buffer at pH 3.0 (CAT#J61391-AK; Alfa Aesar) such that the mixture had an RNA concentration of 0.10 mg/ml. Lipids were stored in ethanol at −20 °C, and RNA constructs were stored in RNAse-free water at −80 °C and were thawed on ice before use. The two phases were prepared at an ethanol:aqueous volume ratio of 1:2, and RNA and lipids combined at an N:P ratio of 5:1. Each phase was loaded into a syringe (BD), and locked onto the NxGen microfluidic cartridge for mixing using a NanoAssemblr Ignite instrument (Precision Nanosystems). The Ignite was set to operate with the following settings: volume ratio- 2:1; flow rate- 12 mL/min; waste volume- 0 mL. The resulting LNPs were then dialyzed into pH 7.4 PBS using Slide-A-Lyzer MINI dialysis devices with a 20K molecular weight cutoff for two rounds at 45 minutes per round.

### LNP characterization

LNPs (2 µg/uL mRNA final concentration) in deionized water were analyzed by dynamic light scattering (DLS) using a Brookhaven Malvern Panalytical DLS system.

For cryoTEM imaging, LNPs were dialyzed in RNAse free water using Slide-A-Lyzer MINI dialysis devices with a 20K molecular weight cutoff for two rounds at 1 hour per round. After dialysis, LNPs were diluted in RNAse water to a concentration of 2 ug/ml. In sample preparation for cryogenic electron microscopy (cryo-EM), 3 μL of the particles sample in buffer containing solution was dropped on a lacey copper grid coated with a continuous carbon film and blotted to remove excess sample without damaging the carbon layer by Gatan Cryo Plunge III. The grid was then mounted on a Gatan 626 single tilt cryo-holder equipped in the TEM column. The specimen and holder tip were cooled down by liquid nitrogen, and the temperature was maintained during transfer into the microscope and subsequent imaging. Imaging on a JEOL 2100 FEG microscope was conducted using a minimum dose method that is essential to avoid sample damage under the electron beam. The microscope was operated at 200 kV and with a magnification in the ranges of 10,000–60,000 for assessing particle size and distribution. All images were recorded on a Gatan 2kx2k UltraScan CCD camera.

### C2C12 cells in vitro transfection

C2C12 murine myoblast cells (ATCC) were cultured in Dulbecco’s Modified Eagle’s Medium (DMEM) containing 10% fetal bovine serum (FBS), and seeded in a 6-well plate at a density of 1x10^6^ cells per well. On the day of transfection, cells were washed 2-3 times in PBS and incubated with 50 µl of mCherry mRNA (concentration 0.1 mg/ml) encapsulated in LNPs or DOTA-LNPs diluted in optim-MEM at 37°C for 4-6 hrs. After 4- 6 hrs, cell media was added on top of treatment and cells were placed back in 37°C for 24 hr. At 24 hours post-transfection, cells were plated in a 96-well U-bottom plate, stained with AquaZombie live/dead stain for 15 minutes at 25°C, and resuspended in flow cytometry buffer then analyzed on a BD FACSCelesta Cell Analyzer.

### Animals and ^64^Cu-labeled LNP immunizations

All animal studies were carried out following an IACUC-approved protocol following state, local, and federal guidelines. DOTA-LNPs were incubated with ^64^Cu (CuCl_2_ in N NaOHI, 30 uCi activity per 10ug mRNA-encapsulated DOTA-LNPs) buffered in pH 7.4 PBS obtained from the Mallinckrodt Institute of Radiology at Washington University School of Medicine at 25°C in a designated radioactivity space. Following 1 hr incubation, loaded LNPs were dialyzed into PBS using Slide-A-Lyzer MINI dialysis devices with a 20K molecular weight cutoff for two rounds at 45 min per round. After dialysis was completed, LNPs were measured for radioactivity (readout in mCi) and loading was validated by instant thin layer chromatography analysis. Eight-week-old BALB/c mice (Jackson Laboratories) were injected intramuscularly in the right gastrocnemius with 10 ug mRNA encapsulated in ^64^Cu -loaded DOTA-LNPs administered in 50 µl PBS. The total radioactivity injected was 12.7-13.7 uCi. Radioactive animals were housed in a separate, designated room, according to Environmental Health & Safety policies, until ten half-lives had elapsed (5.3 d for ^64^Cu).

Indian rhesus macaques were maintained in accordance with the regulations of the Guide for the Care and Use of Laboratory Animal at New Iberia Research Center, University of Louisiana at Lafayette. DOTA-LNPs were incubated with ∼4 mCi ^64^Cu (CuCl_2_ in 0.1N NaOHI) buffered in pH 7.4 PBS obtained from the Department of Medical Physics at the University of Wisconsin in Madison at 37°C in a designated radioactivity space. Following 1 hr incubation, loaded LNPs were dialyzed into PBS using Slide-A-Lyzer MINI dialysis devices with a 20K molecular weight cutoff for two rounds at 45 min per round. After dialysis was completed, LNPs were measured for radioactivity (readout in mCi). A first cohort of 4 animals were injected intramuscularly in the left quadriceps and right deltoid with 25 µg mCherry-encoding mRNA encapsulated in DOTA-LNP and 25 µg mCherry-encoding mRNA encapsulated in diD-labeled LNP in 500 µl PBS. The total radioactivity injected was 3.0 mCi. A second cohort of 4 animals were injected intramuscularly in the right deltoid with µg N332-GT2 trimer-encoding mRNA encapsulated in DOTA-LNP and 25 µg N332-GT2 trimer-encoding mRNA encapsulated in diD-labeled LNP in 500 µl PBS. The total radioactivity injected was 3.0 mCi. Radioactive animals were housed in a separate, designated room, according to EH&S policies, with daily radioactive measures, until ten half-lives had elapsed (5.3 d for ^64^Cu).

### PET-CT imaging of mice and ex vivo tissue biodistribution

Immunized mice were imaged at 0, 3, 6, and 24h post-injection using a G8 PET/CT preclinical small-animal scanner (PerkinElmer). Mice were anesthetized with isoflurane (2% mixed with oxygen) and kept warm using controlled heating pads during the PET/CT scan. Animals were imaged with a static PET scan for 10 minutes followed by a 1.5-minute µCT for anatomical reference. Images were reconstructed with the default 3D maximum likelihood estimation method with CT attenuation correction. At 24 hr, animals were euthanized and tissues were collected for *ex vivo* LNP biodistribution analysis using a Wizard2 automatic gamma counter (PerkinElmer). Measured activity standards were used to calibrate counts to injected dose measurements. Tissues were weighed individually and decay corrected as needed.

### PET-CT imaging of non-human primates

PET/CT scans were performed on a Philips Gemini TF16 scanner. Animals were imaged live at 0, 4, and 24 hr post-injection, and following euthanasia at 40 hr, select tissues were harvested for *ex vivo* imaging. Acquisitions were done in 3D mode with an axial field of view of 57.6 cm.

### PET-CT imaging analysis

PET-CT data was analyzed using the Multi-Image Analysis GUI (MANGO, Research Imaging Institute, UT Health San Antonio) and AMIDE software. Decay-adjusted PET images (corrected to the time of vaccine injection) were normalized by the injected doses and body weight to generate Standardized Uptake Value (SUV) maps. To designate Regions of Interest (ROIs), spherical outlines were placed on target tissues, freehand outline drawings were made, and/or tissue contours on the SUV maps were identified through signal thresholding. These ROIs were then subjected to statistical evaluations to determine SUV max, SUV mean, and SUV sum. The three-dimensional (3D) image data are exhibited as color-coded maximum intensity projections (MIPs) of the SUV max maps.

### NHP tissue extraction, immunofluorescence tissue imaging

Tissue samples were flash frozen, embedded in OCT, and cryosectioned at a thickness of 50 um. Sectioned slides were fixed for 10 min at 25°C with 4% paraformaldehyde, washed with PBS, and perimeters were drawn around samples with a delimiting hydrophobic pen. Samples were treated with Universal Fc receptor blocker (NB309-30) for 30 min at 4°C, followed by goat serum blocking buffer incubation for 60 min at 25°C. Goat anti-human IgD primary antibody (Southern Biotech Cat No. 2030-30) was diluted in blocking buffer and added to tissue samples at 0.02 mg/mL for 24 hr at 4°C. After primary antibody incubation, samples were rinsed 3 times with PBS, and mounted with Prolong Diamond Antifade Mountant (Fisher Scientific Cat No. P36970). Slides were left to cure overnight at 4°C and stored at 4°C until imaged.

Tissue sections were imaged with a 25x water objective or 63x oil objective for immunofluorescence on a Leica SP8 Laser Scanning Confocal Microscope.

### Flow cytometry of lymph nodes

Lymph nodes of vaccinated macaques were isolated 40 hours post-immunization, at the completion of PET imaging. The collected lymph nodes were first scanned before being mechanically dissociated into single cell suspensions. Cells were filtered, counted, and resuspended in freezing medium before storage in liquid nitrogen for subsequent analysis by flow cytometry and single cell RNA sequencing.

For flow cytometry analysis, Zombie UV fixable viability dye (Biolegend) was used according to manufacturer’s protocol. Cells were washed and incubated with Human TruStain FcX (Fc Receptor Blocking Solution) for 15 minutes at 25°C, followed by staining with a cocktail of fluorescent antibodies for 30 minutes at 4°C (panels 1 or 2 described below). After staining cells were fixed with 4% paraformaldehyde for 20 minutes at 4°C. After staining, samples were spiked with Precision Count Beads and cell numbers were calculated according to the manufacturer’s protocol. Samples were acquired on an FACSymphony A3 (BD Biosciences) and data was analyzed using FlowJo v10 (FlowJo Inc).

Panel #1 included anti-human CD16 BUV396 (3G8, BDBiosciences, 1:20), CD14 BUV737 (M5E2, BD Biosciences, 1:20), CD123 BV421 (6H6, Biolegend, 1:20), CD80 BV650 (L307.4, BD Biosciences, 1:20), HLA-DR FITC (Tu36, BDBiosciences, 1:20), CD40 (5C3, BioLegend, 1:10), CD11c PE-Cy7 (3.9, Biolegend, 1:20), CD66abce APC- Vio770 (TET2, Miltenyi, 1:20).

Panel #2 included anti-human CD20 BUV737 (L27, BD Biosciences, 1:20), CD80 BV650 (L307.4, BD Biosciences, 1:20), PD1 BUV785 (EH12.2h7, Biolegend, 1:10), CD4 AF488 (OKT4, Biolegend,1:20), HLA-DR PE (Tu36, Biolegend, 1:20), CXCR5 PE- CY7 (Mu5UBEE, Invitrogen, 1:10), CD8 APC-Cy7 (RPA-T8, Biolegend, 1:50), CD14 APC-Cy7 (M5E2, Biolegend, 1:50), CD16 (3G8, Biolegend, 1:50). For both panels we also collected fluorescent signal for mCherry and the lipophilic dye DiD (APC channel).

### Statistics

All statistical tests and graphical figures were generated using GraphPad Prism Version 10.0.3.

