## Supplementary Materials for "Visualizing lipid nanoparticle trafficking for mRNA vaccine delivery in non-human primates"

Maureen Buckley *et al.*

**This PDF file includes:**

Figs S1 to S8

Table S1

Videos S1 to S10 Captions

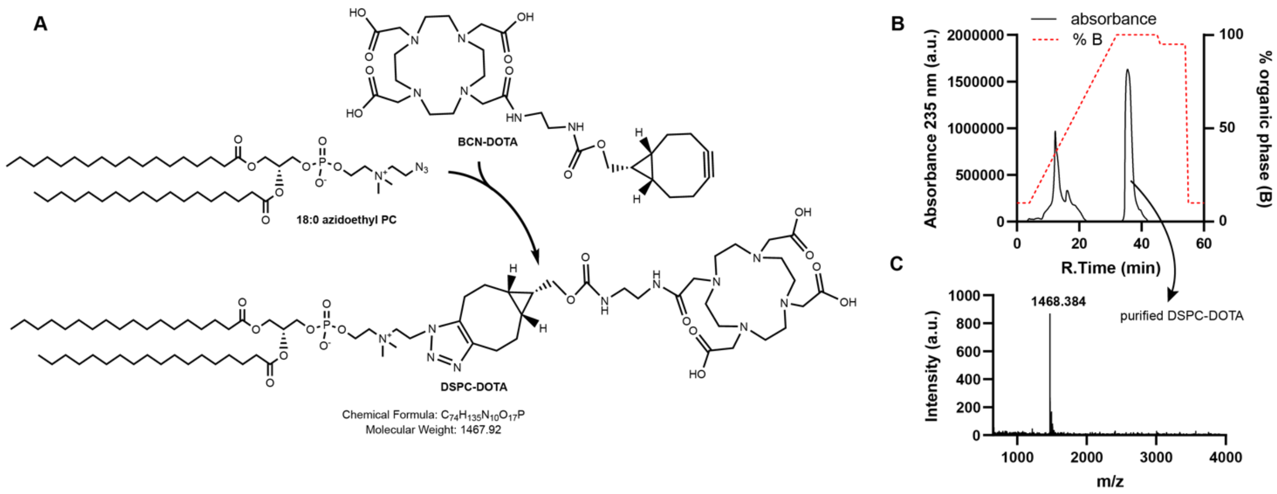


**Figure S1. DSPC-DOTA synthesis and quality validation. (A)** Reaction scheme for DSPC-DOTA synthesis. **(B)** post-reaction purification via RP-HPLC **(C)** final purified product validation with mass spectrometry.

| | | |
|---|---|---|
| **Column** | Jupiter C4 | |
| **Column Temperature** | 40 °C | |
| **Aqueous Phase (A)** | 0.1 M TEAA Water | |
| **Organic Phase (B)** | Methanol | |
| **Flow Rate** | 1 ml/min | |
| **Gradient** | Time | %B |
| | 0 | 10 |
| | 4 | 10 |
| | 32 | 100 |
| | 45 | 100 |
| | 46 | 95 |
| | 54 | 95 |
| | 55 | 10 |
| | 60 | 10 |

**Table S1.RP-HPLC parameters for purification of DSPC-DOTA.**

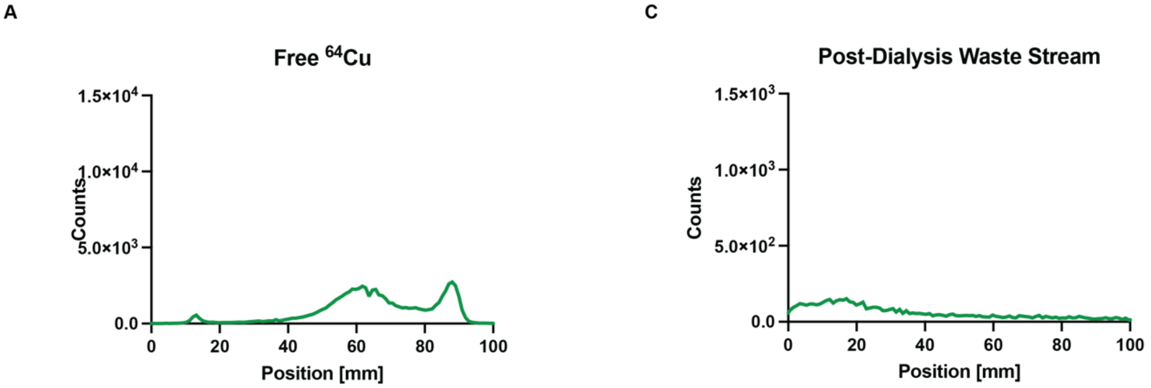


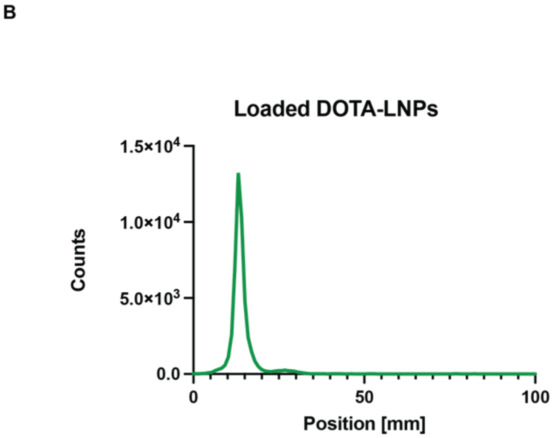


D

| Group | Activity [uCi] |
|---|---|
| DOTA-LNP | 13.7 +/- 0.3 |
| 64Cu Con- | 12.7 +/- 0.6 |

**Figure S2. Validation of $^{64}$Cu loading onto DOTA-LNPs. (A)** Thin layer chromotography analysis for free $^{64}$Cu in PBS. **(B)** Thin layer chromotography analysis for waste steam post-LNP dialysis completed after $^{64}$Cu loading shows minimal trace of free $^{64}$Cu. **(C)** Thin layer chromotography analysis of loaded DOTA-LNPs traces one clean peak that marks successful loading. **(D)** Measured activity in microcuries for DOTA-LNP and free $^{64}$Cu doses injected for PET-CT study.

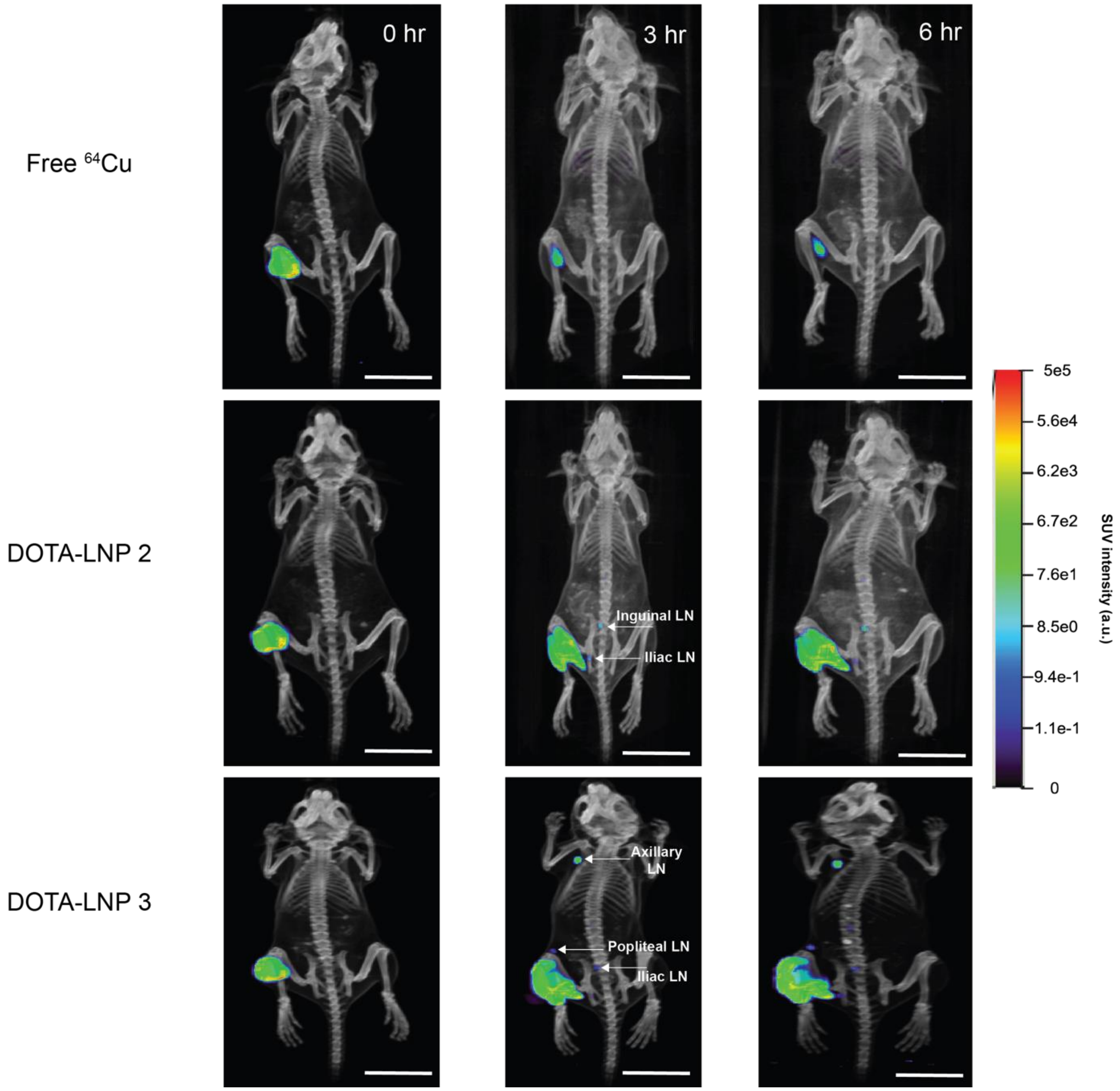


**Figure S3. PET-CT projections for $^{64}$Cu and remaining DOTA-LNP mice.** Projections rendered in AMIDE software with thresholds set constant across all groups/individual mice. Scale bars 50 mm.

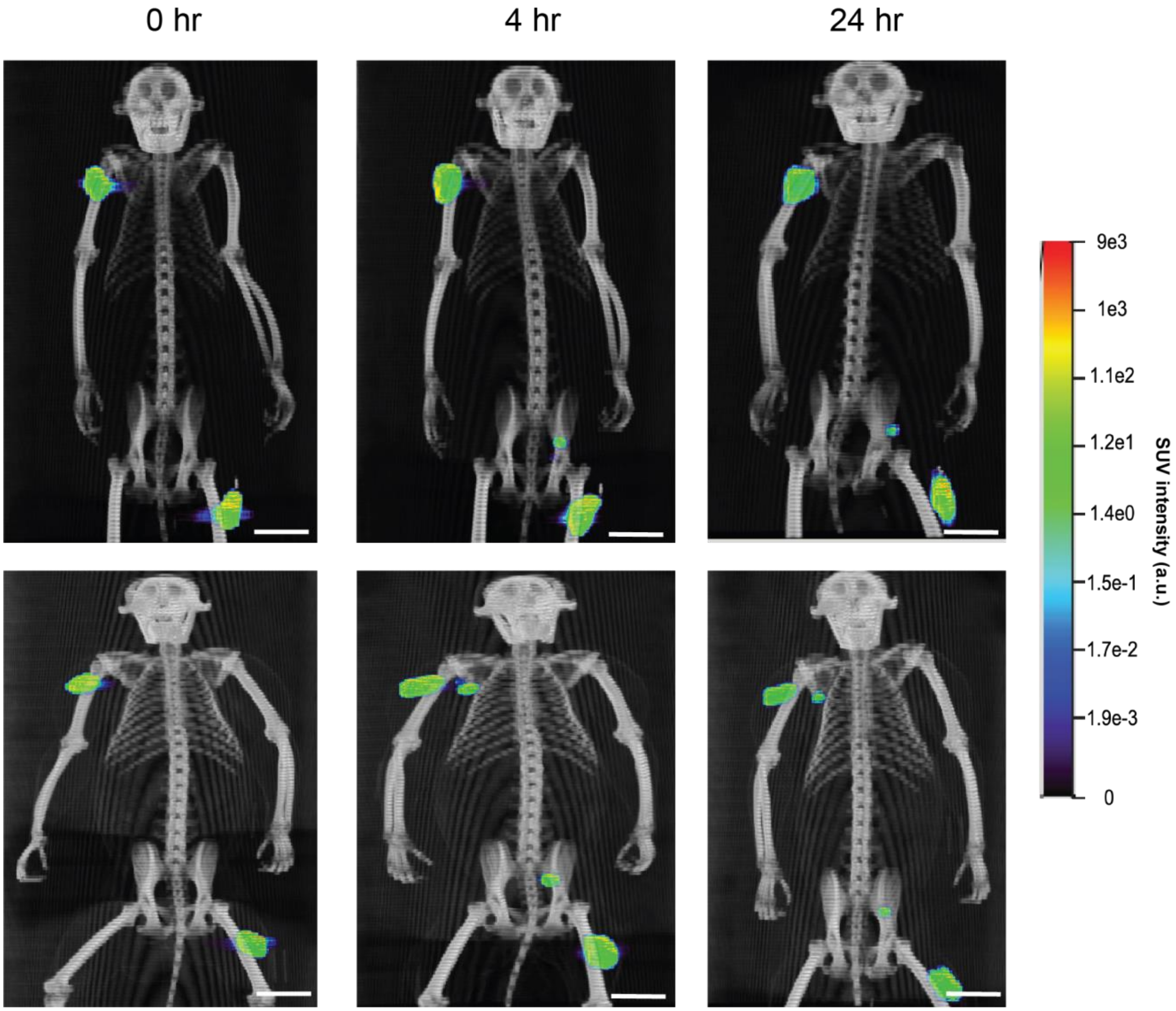


**Figure S4. PET-CT projections for two-injection DOTA-LNP immunized NHPs.** Projections rendered in AMIDE software with thresholds set constant across all animals. Scale bars 50 mm.

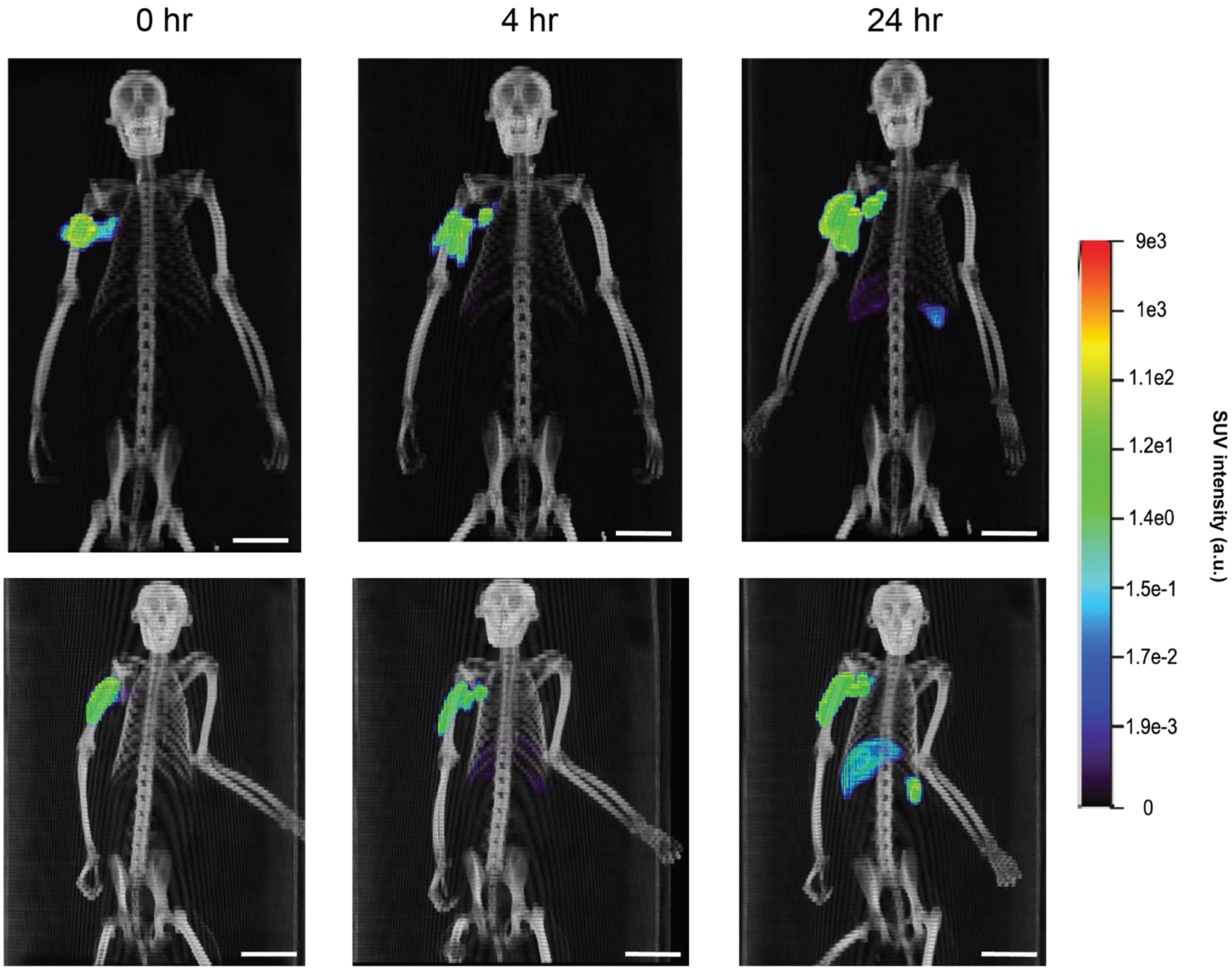


**Figure S5. PET-CT projections for single-injection DOTA-LNP immunized NHPs.** Projections rendered in AMIDE software with thresholds set constant across all animals. Scale bars 50 mm.

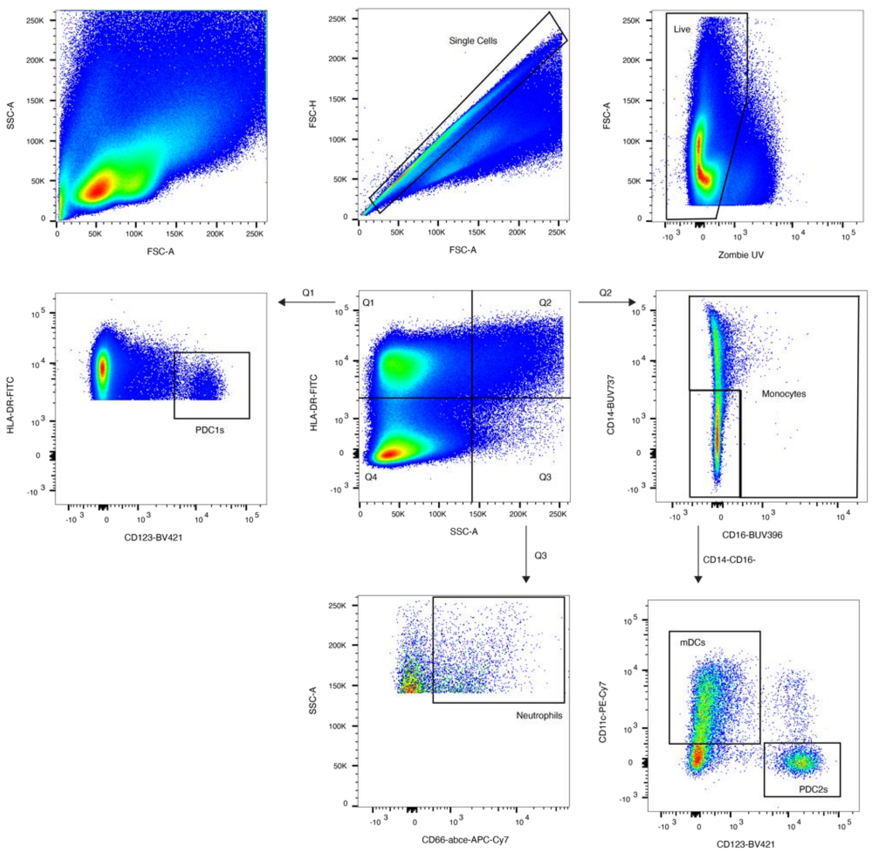


**Figure S6. Flow cytometry gating strategy for myeloid cell analysis.**

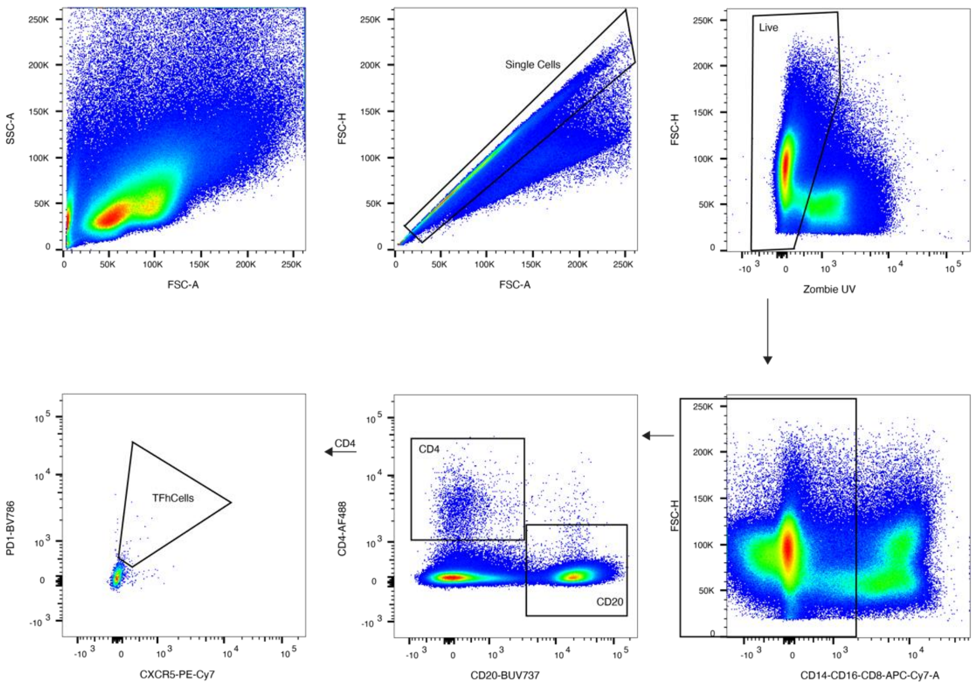


**Figure S7. Flow cytometry gating strategy for lymphocyte analyses.**

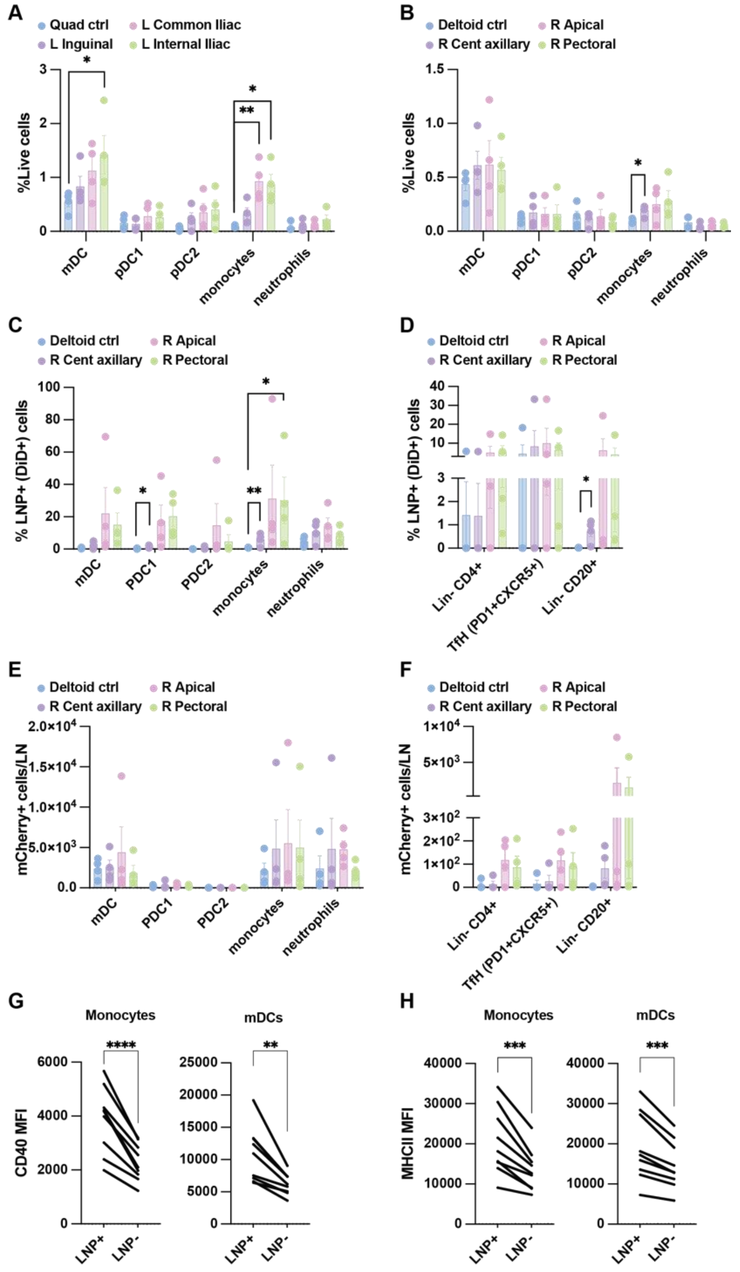


**Figure S8. APCs found to be cell type responsible for LNP uptake and mRNA translation in draining LNs. (A)** Myeloid infiltrate as % of live cells in LNs draining the quadricep injection site. **(B)** Myeloid infiltrate as % of live cells in LNs draining the deltoid injection site. **(C)** Percentage cells positive for LNP signal within specific myeloid populations in LNs draining the deltoid injection site. **(D)** Percentage cells positive for LNP signal within specific lymphoid populations in LNs draining the deltoid injection site. **(E)** Number of cells positive for $mCherry^+$ signal within specific myeloid populations in LNs draining the deltoid injection site. **(F)** Number of cells positive for $mCherry^+$ signal within specific lymphoid populations in LNs draining the deltoid injection site. **(G)** Geometric mean fluorescent intensity of CD80-PerCP-Cy5.5 signal in $LNP^+$ relative to $LNP^-$ cells within LNs with high LNP uptake (>1%). **(H)** Geometric mean fluorescent intensity of MHCII-FITC signal in $LNP^+$ relative to $LNP^-$ cells within LNs with high LNP uptake (>1%). Cell populations were gated as follows: mDCs ($SSC^{high}CD14^-CD16^-$ $MHCII^+CD11c^+$); pDC1s ($SSC^{low}MHCII^+CD123^+$); pDC2s ($SSC^{high}$CD14-CD16-$MHCII^+CD11c^-CD123^+$); Monocytes ($SSC^{high}MHCII^+CD14^+$and/or$CD16^+$); Neutrophils ($SSC^{high}MHCII^-CD66abce^+$); $Lin^-$($CD8^-CD14^-CD16^-$

**Supplementary Video Captions**

**Video S1.** Mouse PET-CT video projection for free $^{64}Cu$ control (from left to right) at 0, 3, and 24 hr timepoints.

**Video S2.** Mouse PET-CT video projection at 0 hr timepoint.

**Video S3.** Mouse PET-CT video projection at 3 hr timepoint.

**Video S4.** Mouse PET-CT video projection at 24 hr timepoint.

**Video S5.** NHP mCherry study PET-CT video projection at 0 hr timepoint.

**Video S6.** NHP mCherry study PET-CT video projection at 4 hr timepoint.

**Video S7.** NHP mCherry study PET-CT video projection at 24 hr timepoint.

**Video S8.** NHP N332-GT2 study PET-CT video projection at 0 hr timepoint.

**Video S9.** NHP N332-GT2 study PET-CT video projection at 4 hr timepoint.

**Video S10.** NHP N332-GT2 study PET-CT video projection at 24 hr timepoint.
